# Cox regression increases power to detect genotype-phenotype associations in genomic studies using the electronic health record

**DOI:** 10.1101/599910

**Authors:** Jacob J. Hughey, Seth D. Rhoades, Darwin Y. Fu, Lisa Bastarache, Joshua C. Denny, Qingxia Chen

**Author notes:** To whom all correspondence should be addressed.

## Abstract

**Background:** The growth of DNA biobanks linked to data from electronic health records (EHRs) has enabled the discovery of numerous associations between genomic variants and clinical phenotypes. Nonetheless, although clinical data are generally longitudinal, standard approaches for detecting genotype-phenotype associations in such linked data, notably logistic regression, do not naturally account for the times at which events occur. Here we explored the advantages of quantifying associations using Cox proportional hazards regression, which can account for the age at which a patient first visited the healthcare system (left truncation) and the age at which a patient either last visited the healthcare system or acquired a particular phenotype (right censoring).

**Results:** Using simulated data, we found that, compared to logistic regression, Cox regression had greater power at equivalent Type I error. We then scanned for genotype-phenotype associations using logistic regression and Cox regression on 50 phenotypes derived from the electronic health records of 49 792 genotyped individuals. In terms of effect sizes, the hazard ratios estimated by Cox regression were nearly identical to the odds ratios estimated by logistic regression. Consistent with the findings from our simulations, Cox regression had approximately 10% greater relative sensitivity for detecting known associations from the NHGRI-EBI GWAS Catalog.

**Conclusions:** As longitudinal health-related data continue to grow, Cox regression may improve our ability to identify the genetic basis for a wide range of human phenotypes.

## Background

The growth of DNA biobanks linked to data from electronic health records (EHRs) has enabled the discovery of numerous associations between genomic variants and clinical phenotypes [1]. Two salient characteristics of EHR data are the large number of correlated phenotypes and the longitudinal nature of observations. Although methods have recently been developed to handle the former [2, 3], approaches to make use of the latter in the context of genome-wide or phenome-wide association studies (GWAS or PheWAS) are less common. Cases are typically defined as individuals with evidence of a phenotype at any timepoint in the EHR, and most large-scale analyses to date have employed logistic or linear regression, which do not naturally account for the time at which a particular event occurs or the highly variable length of observation between patients.

Statistical modeling of time-to-event data has been well studied and frequently applied to the clinical domain [4]. One such method often used to identify genotype-phenotype associations is Cox (proportional hazards) regression [5]. Previous work has demonstrated the advantages of Cox regression over logistic regression for data having a small number of single-nucleotide polymorphisms (SNPs) or collected under particular study designs [6, 7]. To our knowledge, the extent to which these findings generalize to analyses of genome-wide, EHR-linked data remains unclear. Unlike most data analyzed by Cox regression, EHR data are collected for the purposes of clinical care and billing, and are only made available secondarily for research. Thus, not only may individuals leave the healthcare system prior to having an event (a common issue known as right censoring), but they enter the system at various ages (a phenomenon called left truncation).

Here we sought to compare the performance of Cox regression and logistic regression for identifying genotype-phenotype associations in genetic data linked to EHR data. Using both simulated and empirical data, we found that Cox regression shows a modest but consistent improvement in statistical power over logistic regression.

## Results

We first compared logistic regression and Cox regression on the basis of their abilities to detect associations in simulated data. Across 1 000 simulations, logistic regression and Cox regression had mean true positive rates of 0.707 and 0.751, respectively (Fig. 1, 95% CI of difference 0. 0419-0.0447, p-value < 2.2·10^-16^ by paired t-test). Both methods had low false negative rates (mean 1.54·10^-6^ for logistic and 2.11·10^-6^ for Cox). Thus, based on our simulations, Cox regression would be expected to detect an additional 4 associations for every 100 true risk alleles, while falsely claiming 0.6 associations for every 10^6^ non-risk alleles.

**Figure 1.**
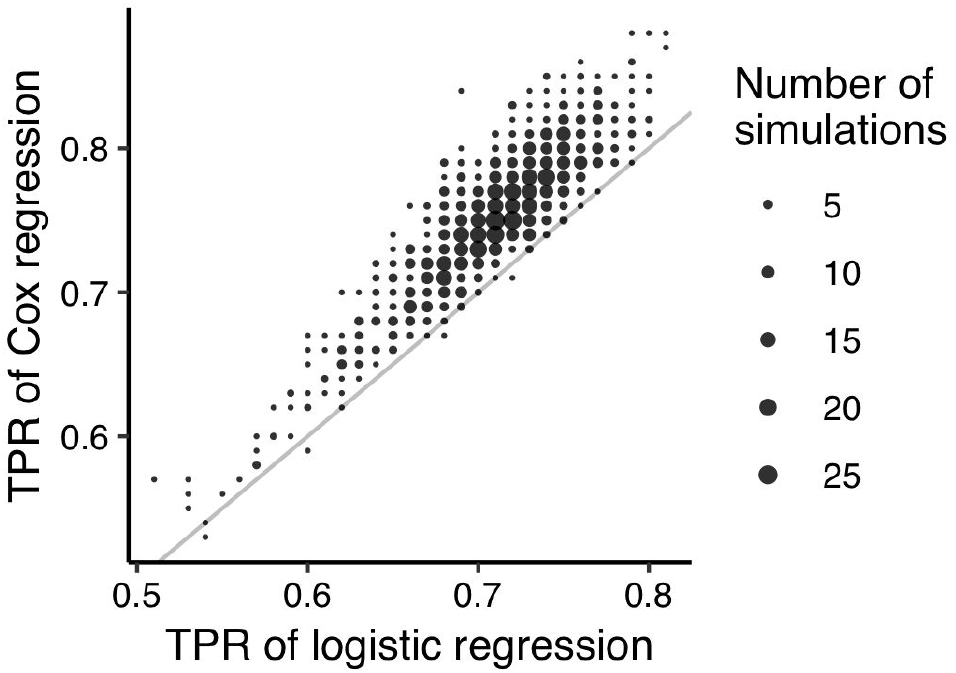
Scatterplot of the true positive rates (TPR) for detecting genotype-phenotype associations in simulated data using logistic regression and Cox regression. Results are based on 1 000 simulations, each of which included 100 risk alleles and 29 900 alleles not associated with the phenotype, and which included left truncation and right censoring. TPR was calculated as the fraction of risk alleles whose having Bonferroni-adjusted p-value < 0.05.

We next compared the two methods using genetic data linked to electronic health records. We selected a cohort of 49 792 individuals of European ancestry, genotyped using the Illumina MEGA platform. We defined 50 phenotypes from the EHR, with the number of cases per phenotype ranging from 104 to 7 972 (Additional file 1: Table S1). For each phenotype, we used Cox regression and logistic regression to run a GWAS on 795 850 common SNPs (including terms for principal components of genetic ancestry, Additional file 2: Figure S1). Overall, the two methods gave similar results (Manhattan plots for four phenotypes in Additional file 2: Figure S2). The p-values were highly correlated and the genomic inflation factors for both methods were generally slightly greater than 1 (Additional file 2: Figure S3A-B). In addition, although coefficients from the two methods have different interpretations, the hazard ratios from Cox regression were nearly identical to the odds ratios from logistic regression (R = 0.9997; Additional file 2: Figure S3C). For associations with a mean −log_10_(P) ≥ 5, however, the p-value from Cox regression tended to be moderately lower than the p-value from logistic regression (Additional file 2: Figure S3D-E). Cox regression also resulted in consistently smaller standard errors of coefficient estimates (Additional file 2: Figure S3F).

We next used the GWAS results from the 50 phenotypes to evaluate each method’s ability to detect known associations from the NHGRI-EBI GWAS Catalog (Additional file 3: Table S2). Across a range of p-value cutoffs, Cox regression had approximately 10% higher relative sensitivity compared to logistic regression (Fig. 2A-B). Even after tailoring the Cox regression procedure for GWAS (see Methods), the implementation of Cox regression we used was 4.5 times slower than logistic regression in PLINK2. A practical strategy to achieve the improved sensitivity without increasing computational cost is to first run logistic regression on all SNPs, then run Cox regression on the subset of SNPs that meet a particular p-value cutoff (Fig. 2A), as previously suggested [7]. The number of hypotheses, and thus the threshold for Bonferroni correction, do not change.

**Figure 2.**
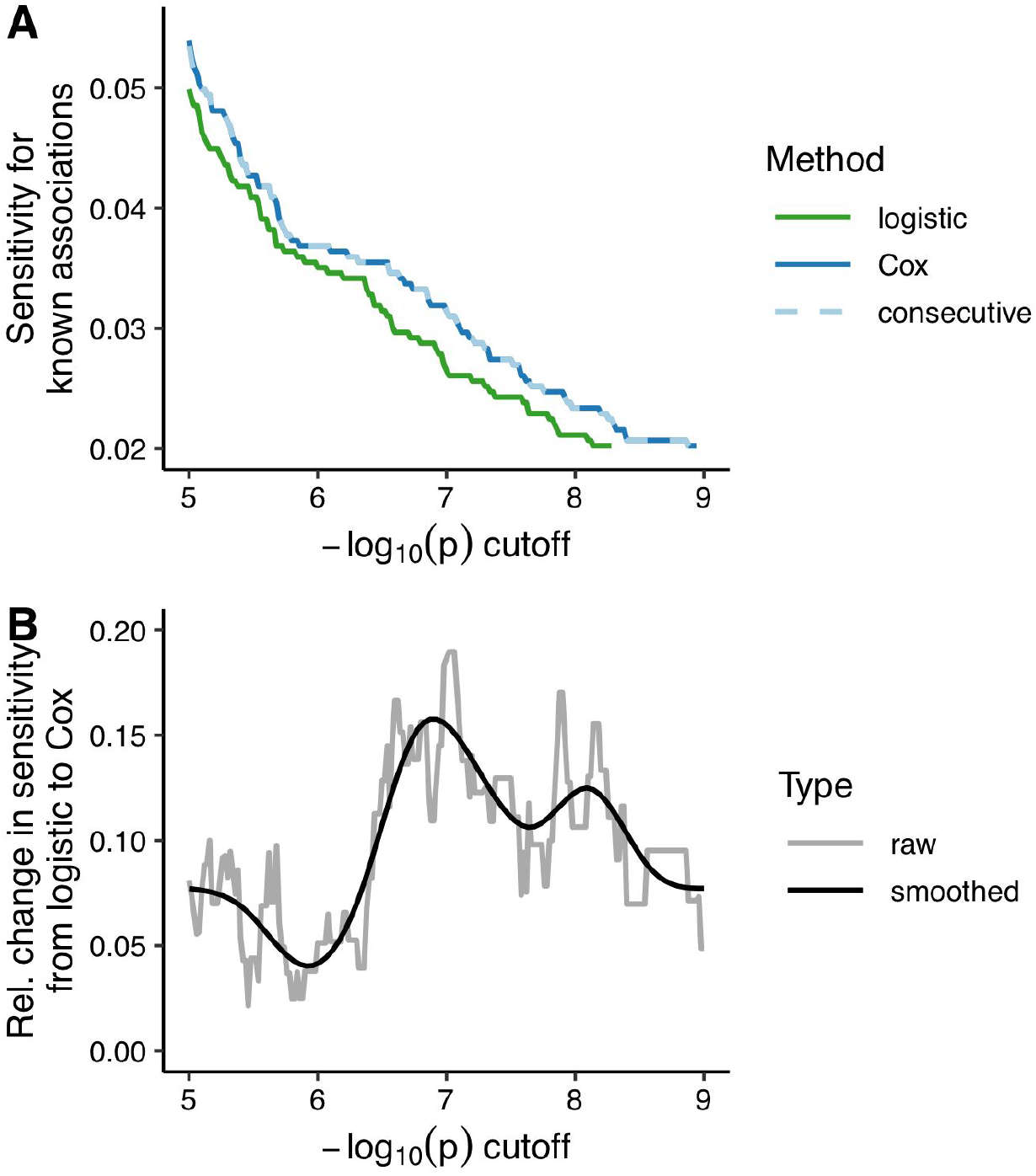
Comparing Cox regression and logistic regression for the ability to detect known genotype-phenotype associations for the 50 phenotypes analyzed. Known significant associations (P ≤ 5·10^-8^) were curated from the NHGRI-EBI GWAS Catalog and aggregated by LD for each phenotype. **(A)** Sensitivity of each method, i.e., fraction of known and tested associations that gave a p-value less than or equal to the specified cutoff. Consecutive refers to the strategy of using the p-value from Cox regression, if the p-value from logistic regression was ≤ 10^-4^. **(B)** Relative change in sensitivity between logistic and Cox regression, i.e., difference between the sensitivities for Cox and logistic, divided by the sensitivity for logistic. The gray line corresponds to the raw value at each cutoff, while the black line corresponds to the smoothed value according to a penalized cubic regression spline in a generalized additive model.

In parallel to quantifying associations using Cox regression, it is natural to visualize them using survival curves. For various phenotype-SNP pairs, we therefore plotted the number of undiagnosed individuals divided by the number at risk (“survival”) as a function of age and genotype (Fig. 3). These curves highlight not only a phenotype’s association with genotype, but also its characteristic age-dependent diagnosis rate.

**Figure 3.**
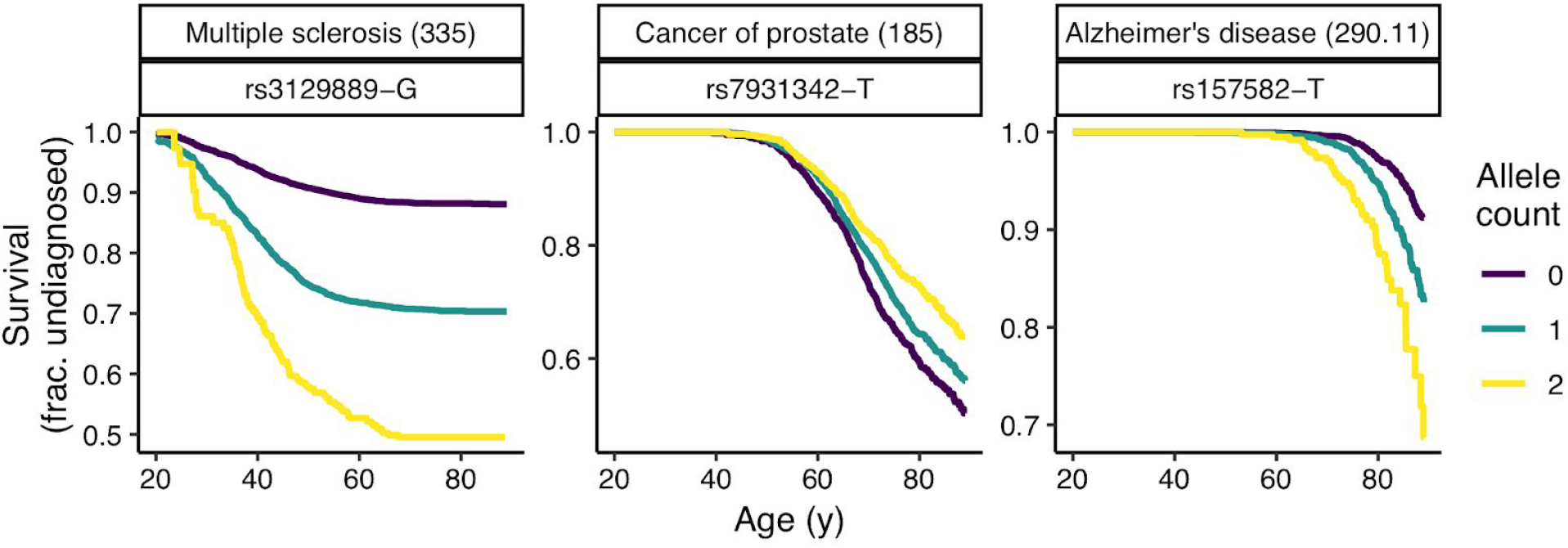
Kaplan-Meier survival curves for three phenotype-SNP pairs. For each phenotype, the corresponding phecode is in parentheses. The curves do not account for sex or principal components of genetic ancestry, and thus are not exactly equivalent to the Cox regression used for the GWAS.

## Discussion

The key piece of additional information required in Cox regression is the time to event. Thus, whereas an odds ratio from logistic regression represents the ratio of cumulative risk over all time, a hazard ratio from Cox regression represents the ratio of instantaneous risk at any given time. In our analysis of EHR data, the time to event corresponded to the age at which a person either received a particular diagnosis code for the second time or was censored. Although acquisition of a diagnosis code is only an approximation for onset of a phenotype, the survival curves for multiple phenotypes suggest that this approximation is valid [8–10].

To account for the fact that most individuals in our data are not observed from birth, we used the age of each individual’s first visit. This formulation of Cox regression, with left truncation and right censoring, corresponds to a counting process [11] and is not currently available in recently published software packages for GWAS of time-to-event outcomes [12, 13]. Furthermore, Cox regression is not available at all in popular GWAS tools such as PLINK. Thus, the implementation of Cox regression we used was not optimized for GWAS. Future work should make it possible to reduce the differences in computational cost and ease of use between Cox regression and logistic regression. In the meantime, we recommend the strategy of logistic followed by Cox [7]. Although the initial threshold for logistic regression is arbitrary, our results suggest that a relatively loose threshold (e.g., P ≤ 10-4) is likely to catch all significant associations while only adding a small amount of computational cost.

Our use of the GWAS Catalog has multiple limitations. First, both methods showed low sensitivity, likely because for half of the 50 phenotypes, the number of EHR-derived cases was in the hundreds, whereas the number of cases from GWAS Catalog studies for these phenotypes was in the thousands. Second, the majority of studies in the GWAS Catalog followed a case-control design and quantified associations using either logistic or linear regression, not Cox regression. Thus, although the GWAS Catalog is the closest we have to a gold standard, it was important that our analyses of simulated data and empirical data gave consistent results.

## Conclusions

Here we used Cox regression to model the time to a single event, i.e., diagnosis of a particular phenotype. In the future, more sophisticated models may be able to account for subsequent response to treatment or semi-continuous traits such as lab values. We are especially interested in the potential of Cox mixed models, which, like linear mixed models [14], use random effects to account for genetic relatedness, an increasingly important factor in EHR-linked samples [15]. Such an approach applied to large-scale datasets such as from the Million Veterans Program or the All of Us Research Program [16, 17], if appropriately adjusted for environmental and societal factors, may enable the creation of clinically useful polygenic hazard scores. Overall, as longitudinal, health-related data continue to grow, accounting for time through methods such as Cox regression may improve our ability to identify the genetic basis for human phenotypes.

## Methods

### Simulation Studies

We compared logistic regression and Cox regression in comprehensive simulations with 1 000 replicates. Simulation parameters were selected to balance similarity to empirical data and computational cost. As the effect sizes estimated by the two methods are not equivalent (i.e., odds ratio versus hazard ratio), we evaluated the methods in terms of average power and type I error calculated from true and false associations in each simulation. The simulations were designed to roughly mimic the empirical study on EHR data. In each simulation, we sampled minor allele counts for 30 000 SNPs in 29 000 individuals from a binomial distribution, with each minor allele’s probability independently simulated from the distribution of minor allele frequencies in the empirical genotype data. For simplicity, we simulated a haploid genome, i.e., each individual had only one allele at each SNP. Of the 30 000 minor alleles, 100 were declared as true risk alleles, with their coefficients sampled from Unif(0.3, 0.5), i.e., a uniform distribution between 0.3 and 0.5. The remaining 29 900 minor alleles were declared as false risk alleles by setting their coefficients to 0. The true event time was simulated from a multivariable Cox regression with baseline hazard generated from Exponential(*λ*) with *λ*=10 000 and the parametric component including all SNPs. The censoring time was simulated from Gamma(1,1) and truncated by 2. The right censored observed event time was the minimum of the true event time and the censoring time. The left truncation time was simulated from Unif(0, 0.1). Individuals whose censoring time or event time was less than the truncation time were removed from the dataset (mean 10% of individuals, range 5.35%-22.7%). The mean event rate was 30.3% (range 8.50%-56.0%). In each simulation, univariate Cox regression (accounting for left truncation) and univariate logistic regression were run for each SNP. Statistical significance was based on Bonferroni correction with an overall type I error rate of 0.05.

### Processing the genotype data

Our empirical data came from the Vanderbilt Synthetic Derivative (a database of de-identified electronic health records) and BioVU (a DNA biobank linked to the Synthetic Derivative) [18]. We used a cohort that was genotyped using the Illumina MEGA platform. To identify individuals of European ancestry (the majority in BioVU), we used STRUCTURE to create three clusters, keeping those individuals who had a score ≥ 0.9 for the cluster that corresponded to European ancestry [19]. We then filtered SNPs to keep those that had a minor allele frequency ≥ 0.01, call rate ≥ 0.95, p-value of Hardy-Weinberg equilibrium ≥ 0.001, and p-value of association with batch ≥ 10^-5^. To calculate the principal components of genetic ancestry, we followed the recommended procedure of the SNPRelate R package v1.16.0 [20]. Specifically, we pruned SNPs based on an LD threshold r = 0.2, then used the randomized algorithm to calculate the first 10 PCs [21].

### Identifying phenotypes for empirical study

To compare the ability of Cox and logistic regression to detect known associations, we selected 50 phenotypes that could be studied with EHR data and which also had known associations from the NHGRI-EBI GWAS Catalog v1.0.2 r2018-08-30 (Additional file 1: Table S1) [22]. The phenotypes were selected before the analysis was performed. We only considered GWAS Catalog studies with at least 1 000 cases and 1 000 controls of European ancestry (Additional file 3: Table S2). We manually mapped studies and their corresponding traits to EHR phenotypes using phecodes, which are derived from billing codes [23]. For each phenotype, we defined cases as individuals who received the corresponding phecode on two distinct dates, and controls as individuals who have never received the corresponding phecode. Each phenotype had at least 100 cases.

### Running the GWAS

For both Cox regression and logistic regression, the linear model included terms for genotype (assuming an additive effect) and the first four principal components of genetic ancestry (Additional file 2: Figure S1). Depending on the phenotype, the model either included a term for biological sex or the cases and controls were limited to only females or only males. For logistic regression, the model also included terms for age at time of last visit (modeled as a cubic smoothing spline with three knots) and length of time between first visit and last visit. For Cox regression, the model used the counting process formulation, such that time 1 corresponded to age at first visit ever and time 2 corresponded to age on the second distinct date of receiving the given phecode (for cases) or age at last visit (for controls).

Logistic regression was run using PLINK v2.00a2LM 64-bit Intel (30 Aug 2018) [24]. Cox regression was run in R using the agreg.fit function of the survival package v2.43-1. The agreg.fit function is normally called internally by the coxph function, but calling agreg.fit directly is faster. The total runtimes for the GWASes of the 50 phenotypes using logistic and Cox regression (parallelized on 36 cores) were 1.6 days and 7.1 days, respectively.

### Comparing GWAS results using the GWAS Catalog

For each mapped study from the GWAS Catalog, we only considered SNPs having an association P ≤ 5·10^-8^. For each phenotype, we then used LDlink [25] to group the associated SNPs into LD blocks (r^2^ ≥ 0.8). For each associated SNP for each phenotype, we then determined which SNPs on the MEGA platform were in LD with that SNP (r^2^ ≥ 0.8), and assigned those SNPs to the corresponding phenotype and LD block. Using the EHR-based GWAS results, we then calculated the sensitivity of Cox regression and logistic regression based on the number of phenotype-LD block pairs for which at least one SNP in that LD block had a p-value less than a given p-value cutoff (across a range of cutoffs).

## Declarations

### Acknowledgements

Not applicable.

### Funding

This work was supported by the U.S. National Institutes of Health (R35GM124685 to JH, T32HG008341 to SR, R01LM016085 to JD, and U24CA194215-01A1 to QC) and the Kleberg Foundation (to JD). The Vanderbilt Synthetic Derivative and BioVU are supported by institutional funding and by CTSA award UL1TR002243 from NCATS/NIH.

### Availability of data and materials

Acces to individual-level EHR and genotype data is restricted by the IRB. Code and summary-level results are available at https://figshare.com/s/0fa5f9549218dc3f02d5.

### Authors’ contributions

JH, SR, LB, and QC designed the study. JH and QC performed the analyses. JH, DF, and QC drafted the manuscript. All authors interpreted the results, edited the manuscript, and read and approved the final manuscript.

### Ethics approval and consent to participate

The Vanderbilt Institutional Review Board reviewed and approved this study as non-human subjects research (IRB# 081418).

### Consent for publication

Not applicable.

### Competing interests

The authors declare they have no competing interests.

### Additional files

Additional file 1: Table S1, Information for each of the 50 phenotypes.

Additional file 2: Figures S1-S3, Supplemental figures for principal components of genetic ancestry and GWAS results using Cox and logistic regression.

Additional file 3: Table S2, Mapping between phecodes and GWAS Catalog study accessions.

