## Supplemental Figures for "Cox regression increases power to detect genotype-phenotype associations in genomic studies using the electronic health record"

Figure S1

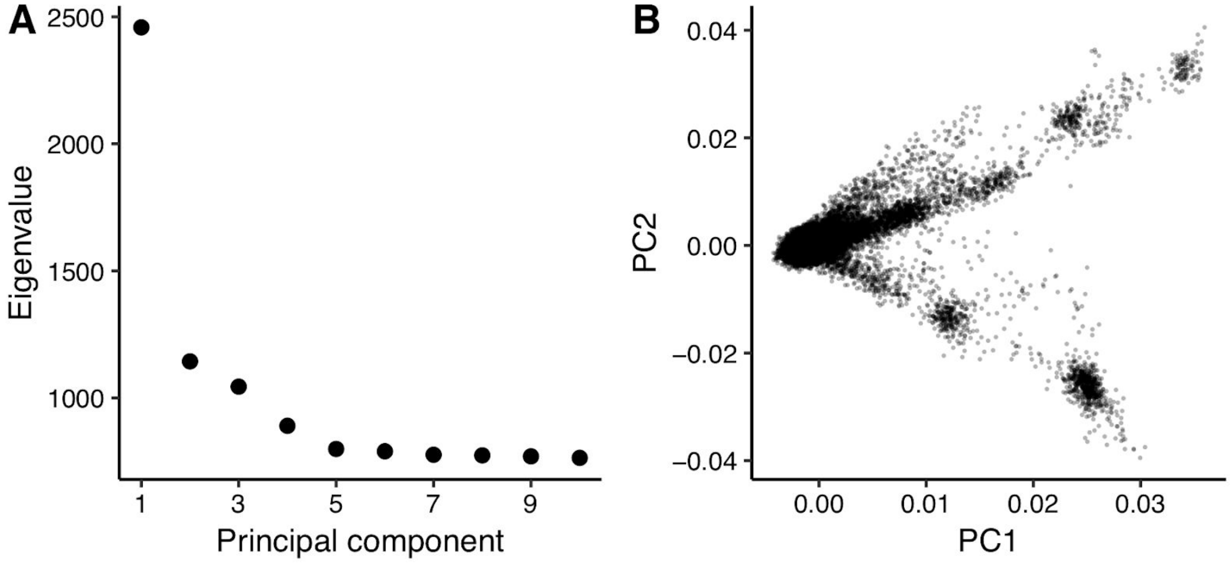


Principal components of genetic ancestry, calculated using SNPRelate. **(A)** Eigenvalues for the top 10 principal components. **(B)** Scatterplot of PC2 vs. PC1, where each point corresponds to a subject in the cohort.

Figure S2

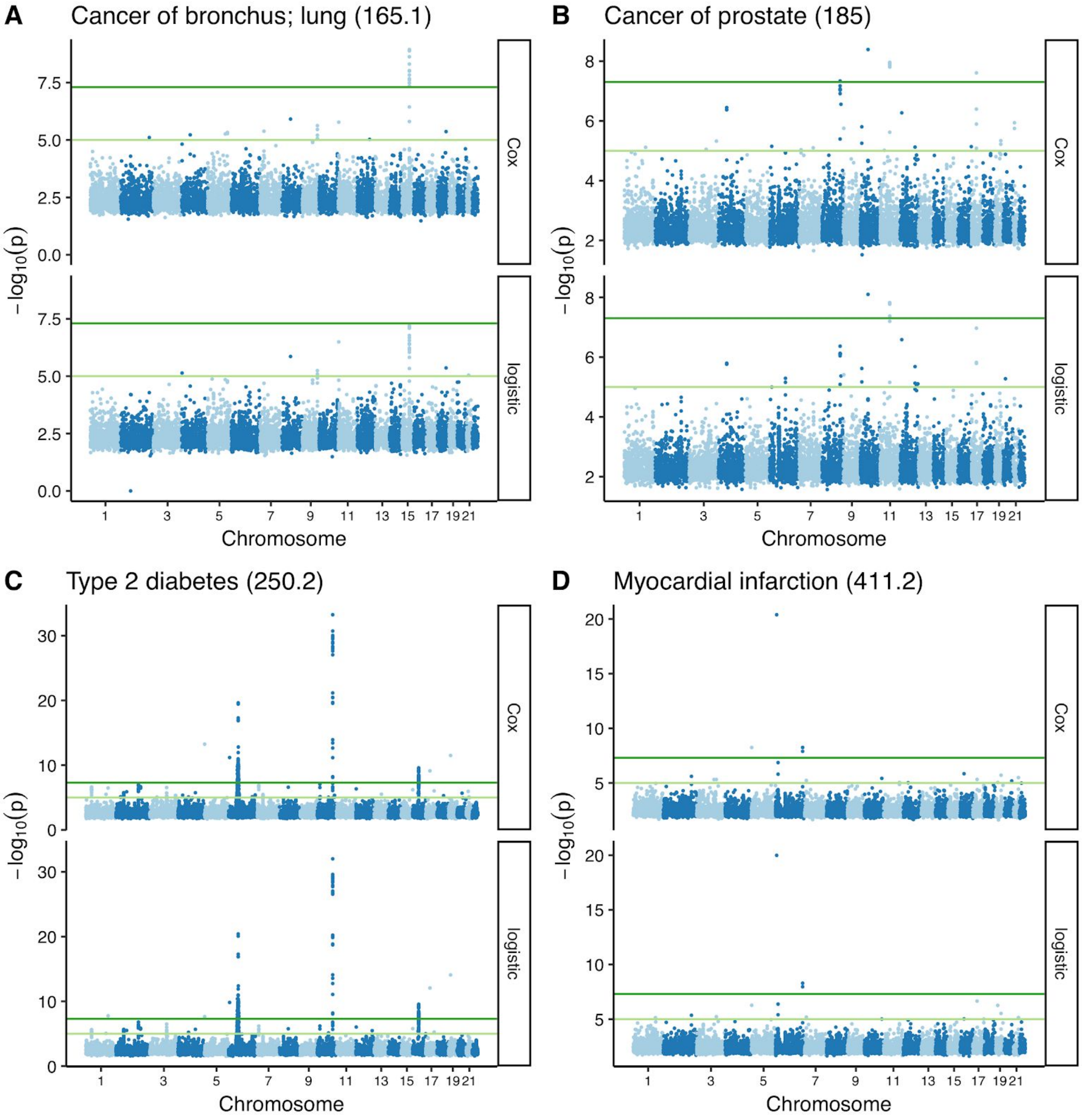


Manhattan plots of GWAS results using Cox and logistic regression for four phenotypes (phecode in parentheses). For each phenotype, only associations having mean($-\log_{10}(P)$) ≥ 2 are shown. Dark green lines correspond to $P = 5 \cdot 10^{-8}$ and light green lines correspond to $P = 10^{-5}$.

Figure S3

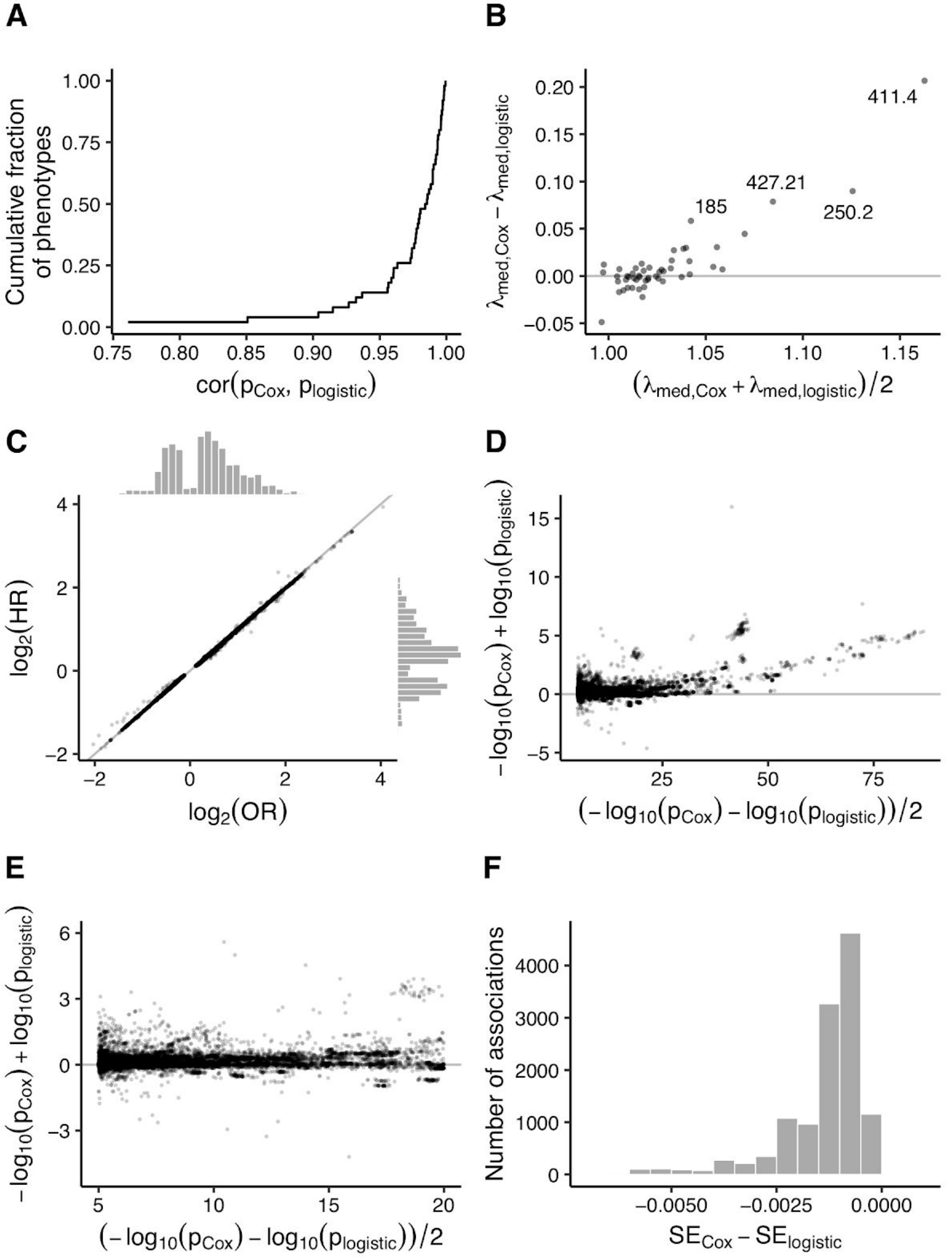
A
Cumulative fraction of phenotypes
1.00
0.75
0.50
0.25
0.00
0.75 0.80 0.85 0.90 0.95 1.00
$cor(p_{Cox}, p_{logistic})$
B
$\lambda_{med,Cox} - \lambda_{med,logistic}$
0.20
0.15
0.10
0.05
0.00
−0.05
411.4
427.21
185
250.2
1.00 1.05 1.10 1.15
$(\lambda_{med,Cox} + \lambda_{med,logistic})/2$
C
$log_2(HR)$
4
2
0
−2
−2 0 2 4
$log_2(OR)$
D
$-log_{10}(p_{Cox}) + log_{10}(p_{logistic})$
15
10
5
0
−5
25 50 75
$(-log_{10}(p_{Cox}) - log_{10}(p_{logistic}))/2$
E
$-log_{10}(p_{Cox}) + log_{10}(p_{logistic})$
6
3
0
−3
5 10 15 20
$(-log_{10}(p_{Cox}) - log_{10}(p_{logistic}))/2$
F
Number of associations
4000
3000
2000
1000
0
−0.0050 −0.0025 0.0000
$SE_{Cox} - SE_{logistic}$

Comparing the results of GWAS based on Cox regression or logistic regression for 50 selected phenotypes. **(A)** Empirical cumulative distribution function of the Pearson correlation between p-values for each phenotype. **(B)** Mean-difference plot of the genomic inflation factor. Each point corresponds to a phenotype. Phenotypes with a difference > 0.05 are labeled with the corresponding phecode (411.4: coronary atherosclerosis, 250.2: type 2 diabetes, 427.21: atrial fibrillation, 185: prostate cancer). **(C)** Scatterplot and marginal histograms of $\log_2$(effect size) for Cox and logistic regression, calculated as $log_2(e^{\beta})$, where $\beta$ is the coefficient for genotype. HR refers to hazard ratio from Cox regression, OR refers to odds ratio from logistic regression. In panels C-E, each point corresponds to a SNP-phenotype pair. Panels C-F include only those associations for which $P \leq 10^{-5}$ for either Cox or logistic regression. **(D)** and **(E)** Mean-difference plots of $-\log_{10}(P)$ with different x- and y-axes. **(F)** Histogram of the difference in standard error of the coefficient estimate.
